# CoolMPS^™^: Advanced massively parallel sequencing using antibodies specific to each natural nucleobase

**DOI:** 10.1101/2020.02.19.953307

**Authors:** Snezana Drmanac, Matthew Callow, Linsu Chen, Ping Zhou, Leon Eckhardt, Chongjun Xu, Meihua Gong, Scott Gablenz, Jyothi Rajagopal, Qing Yang, Christian Villarosa, Anthony Au, Kyle Davis, Alexander Jorjorian, Jingjing Wang, Ao Chen, Xian Zhang, Adam Borcherding, Xiaofang Wei, Mingxuan Zhang, Yonghui Xie, Nina Barua, Jay Shafto, Yuliang Dong, Yue Zheng, Lin Wang, Lili Zhai, Jiguang Li, Sha Liao, Wenwei Zhang, Jian Liu, Hui Jiang, Jian Wang, Handong Li, Xun Xu, Radoje Drmanac

## Abstract

Massively parallel sequencing (MPS) on DNA nanoarrays provides billions of reads at relatively low cost and enables a multitude of genomic applications. Further improvement in read length, sequence quality and cost reduction will enable more affordable and accurate comprehensive health monitoring tests. Currently the most efficient MPS uses dye-labeled reversibly terminated nucleotides (RTs) that are expensive to make and challenging to incorporate. Furthermore, a part of the dye-linker (scar) remains on the nucleobase after cleavage and interferes with subsequent sequencing cycles. We describe here the development of a novel MPS chemistry (CoolMPS™) utilizing unlabeled RTs and four natural nucleobase-specific fluorescently labeled antibodies with fast (30 sec) binding. We implemented CoolMPS™ on MGI’s PCR-free DNBSEQ MPS platform using arrays of 200nm DNA nanoballs (DNBs) generated by rolling circle replication and demonstrate 3-fold improvement in signal intensity and elimination of scar interference. Single-end 100-400 base and pair-end 2×150 base reads with high quality were readily generated with low out-of-phase incorporation. Furthermore, DNBs with less than 50 template copies were successfully sequenced by strong-signal CoolMPS™ with 3-times higher accuracy than in standard MPS. CoolMPS™ chemistry based on natural nucleobases has potential to provide longer, more accurate and less expensive MPS reads, including highly accurate “4-color sequencing” on the most efficient dye-crosstalk-free 2-color imagers with an estimated sequencing error rate of 0.00058% (one error in 170,000 base calls) in a proof-of-concept demonstration.

## Introduction of Massively Parallel Sequencing (MPS)

MPS (*1*–*6*) is driving advanced genomics applications (*7*–*9*) by providing billions of sequence reads from patterned DNA nanoarrays (*4*). Longer, paired-end and barcoded (stLFR) reads (*10*) are enabling broader applications. Further improvement in read length, quality and cost reduction will enable more challenging sequencing-based health monitoring tests that need to be comprehensive, accurate and affordable. Furthermore, a full understanding of our human genetic program will require genome sequencing for millions if not billions of people and deep molecular characterization of millions of our cells by various single cell “omics” tests requiring extremely large scale sequencing. Currently, most advanced MPS methodologies are comprised of sequencing cycles incorporating labeled reversible terminated nucleotides (RTs) proposed almost 30 years ago (*11*). Ion Torrent (*5*) uses natural nucleotides (one at a time) that can result in higher errors in homopolymers. Base-labeled RTs have several limitations including efficiency of incorporation, cost of synthesis, single dye per base limited signal, and incomplete regeneration of natural nucleotides (part of the dye linker is left on the base after dye cleavage as a “scar”). We describe here the development of a novel, potentially more efficient and accurate MPS chemistry using a pool of all four unlabeled RTs with natural nucleobases.

## CoolMPS™: a new MPS chemistry using unlabeled RTS

A unique and distinguishing feature of the CoolMPS™ chemistry is that no fluorescently labeled RTs are required (Fig. 1). CoolMPS™ chemistry was proposed by one of us (R. Drmanac) in 2016 and described in a patent application (*12*). Incorporation of unlabeled reversibly terminated nucleotides, and base determination, is performed in each cycle of sequencing using base-specific 3’ block-dependent fluorescently labeled antibodies. Removal of the bound antibodies and 3’ blocking moiety on the sugar group of the nucleotide regenerates natural nucleotides with no scar on the base. This feature of reversion to a natural nucleotide allows further extension of the strand in a new cycle of sequencing without any interference from the prior cycle. Furthermore, unlabeled RTs are easier and less costly to make, and they can be incorporated more efficiently. An additional advantage of CoolMPS™ is that antibodies can carry multiple molecules (e.g. 2-5) of the same dye, greatly increasing sequencing signal compared with single dye per base on standard labeled RTs.

**Figure 1:**
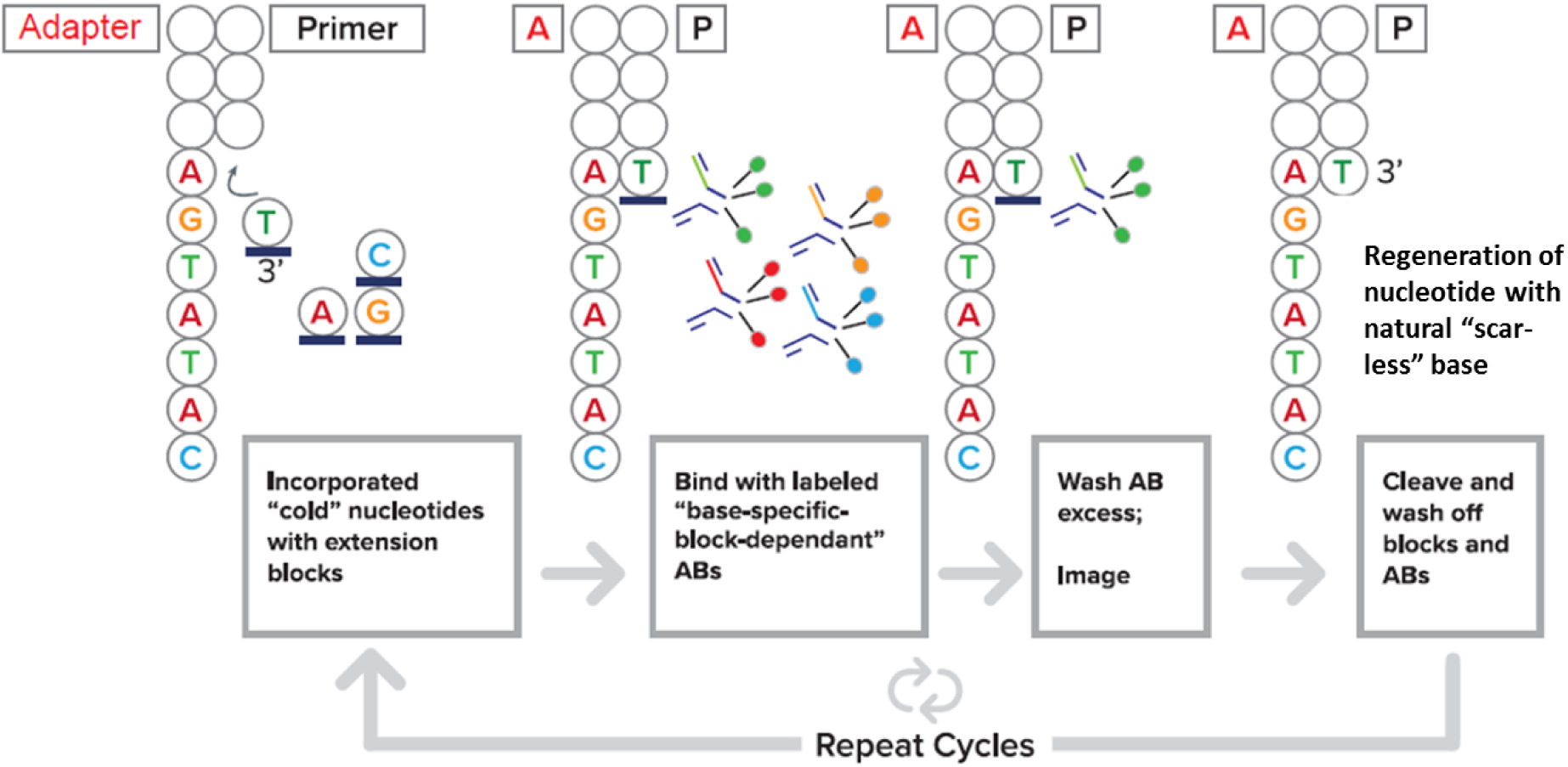
CoolMPS™ process overview. Bars (‐) on the unlabeled (“cold”) nucleotides depict removable 3’ chemical blocks. Antibodies specific for RTs with natural nucleobase are depicted with three dye molecules to increase fluorescent signal.

## First CoolMPS™ implementation

### Obtaining and labeling monoclonal CoolMPS™ antibodies

Immunization: To demonstrate CoolMPS™ we used natural unlabeled adenosine, cytosine, guanosine and thymidine, monophosphate nucleotides with a 3’-O-azidomethyl blocking group (Fig. 2). This cleavable blocking group was originally synthesized by Zavgorodny in 1991 who highlighted its triggerable cleavage in mild conditions (*13*). The group is relatively small enabling easier incorporation by DNA polymerases, is stable during RT incorporation and imaging, and allows fast cleavage by a well-known chemical reaction. The stability and small size of this 3’ blocking group relative to natural nucleobases also improves the chances of obtaining nucleobase specific antibodies with binding equally or more dependent on the nucleobase than on the 3’ blocking group. To generate needed antibodies, the blocked nucleotides were linked via the monophosphate to an N-hydroxysuccinimide group (Fig. 2) which was then linked to KLH protein for the immunization of rabbits every two weeks. Immunogens were custom made by AAT Bioquest (Sunnyvale, CA) and monoclonal antibodies were custom produced by Yurogen Biosystems LLC (Worcester, MA). Sera was collected from immunized rabbits over a three-month period and screened by ELISA to determine immune response. Antigen for the ELISA screen was 3’-blocked nucleotides linked to BSA coated onto wells of a microtiter plate.

**Figure 2:**
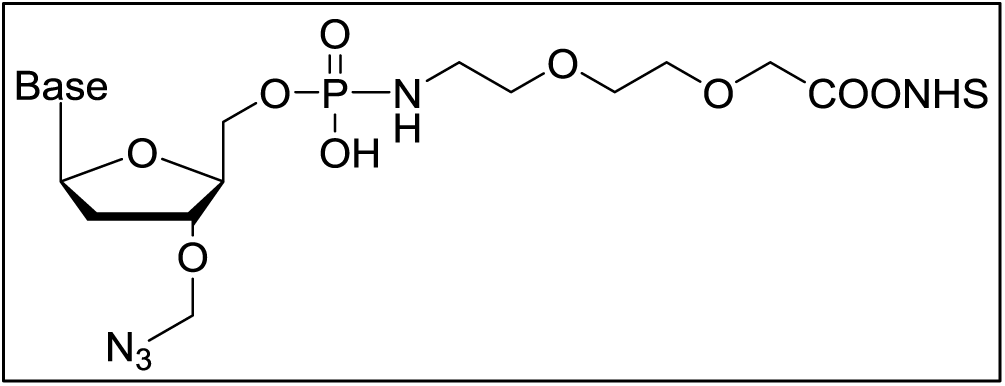
Example structure of the immunogen used to generate rabbit monoclonal antibodies. The NHS ester was first reacted with the KLH protein before immunization.

Splenocyte screening: Splenocytes collected from sero-positive rabbits were FACS sorted for positive antibody expression using antigen bound via biotin to fluorescently labeled streptavidin. FACS selected single cells with positive expression for immunogen reactive surface bound IgG for further growth in 384-well plates. This allowed confirmative screening of expressed antibodies.

Antibody screening: After splenocyte expansion, supernatant from each single cell derived clonal culture was screened against all 4 nucleotide variants (A, C, G and T) to identify clones giving high reactivity against the specific nucleobase antigen, and low or non-detectable reactivity to the 3 non-targeted bases. Those antibodies with high non-specific binding (>20%), as indicated by high ELISA positive signal to the non-targeted bases, were excluded from further consideration.

Antibody cloning and expression: Selected splenocyte cultures had coding regions for antibody heavy and light chains cloned into a plasmid expression system. These plasmids were used to transiently transfect a 293 cell-line for monoclonal antibody production. Expressed antibodies were purified by protein A capture columns and eluted in low pH buffer before buffer exchange into phosphate buffered saline.

Antibody labeling: Antibodies were labeled by reaction of available free amines on the protein with NHS ester activated fluorescent dyes (*14*). NHS ester activated fluorophores were diluted in anhydrous DMSO and reacted at concentrations (10-100 μM) that provide strong signals without adversely affecting antibody binding or specificity. Relatively low and easy to obtain concentrations of antibody (1 mg/ml) were adjusted to pH 8 in bicarbonate buffer and reacted with the NHS ester dyes. Incubation was continued for 45 min at room temperature before quenching of unreacted dye in tris-buffered saline (pH 7.4). Without any purification, these labeled antibodies were aliquoted and stored at -20C.

### Characterization of CoolMPS™ antibodies in sequencing assays

#### Sequencing platform

DNBSEQ-G400, MGI’s MPS platform (MGI is a BGI subsidiary in Shenzhen, China), was used for testing and implementing the CoolMPS™ process. The DNBSEQ platform utilizes PCR-free nanoarrays of DNA nanoballs (DNBs); linear concatemers of DNA copies generated by rolling circle replication that are bound to defined positions of a patterned nanoarray (*4*). Amplification occurs through continuous copying of the original circular DNA template, avoiding creation of clonal errors that are unavoidable in PCR amplification.

For popular pair-end (PE) or second-end sequencing on the DNBSEQ platform, we recently developed a controlled multiple displacement amplification (MDA) process on DNB arrays (*15*). After the first read is generated on DNBs, extended products (optionally using an additional primer) are further extended using natural unblocked nucleotides in a controlled and sufficiently synchronized way by a strand displacement polymerase such as Phi29. The process generates single-stranded (ss) DNA branches complementary to original DNBs and still bound to DNBs through regions that are not displaced (Fig. 3) The resulting “branched DNBs” usually comprise 1-3 template copies per branch, providing more priming sites and stronger signal in the second end-read than in the first end-read. Current commercial MGI pair-end kits for DNBSEQ platform include PE100, PE150 and PE200.

**Figure 3:**
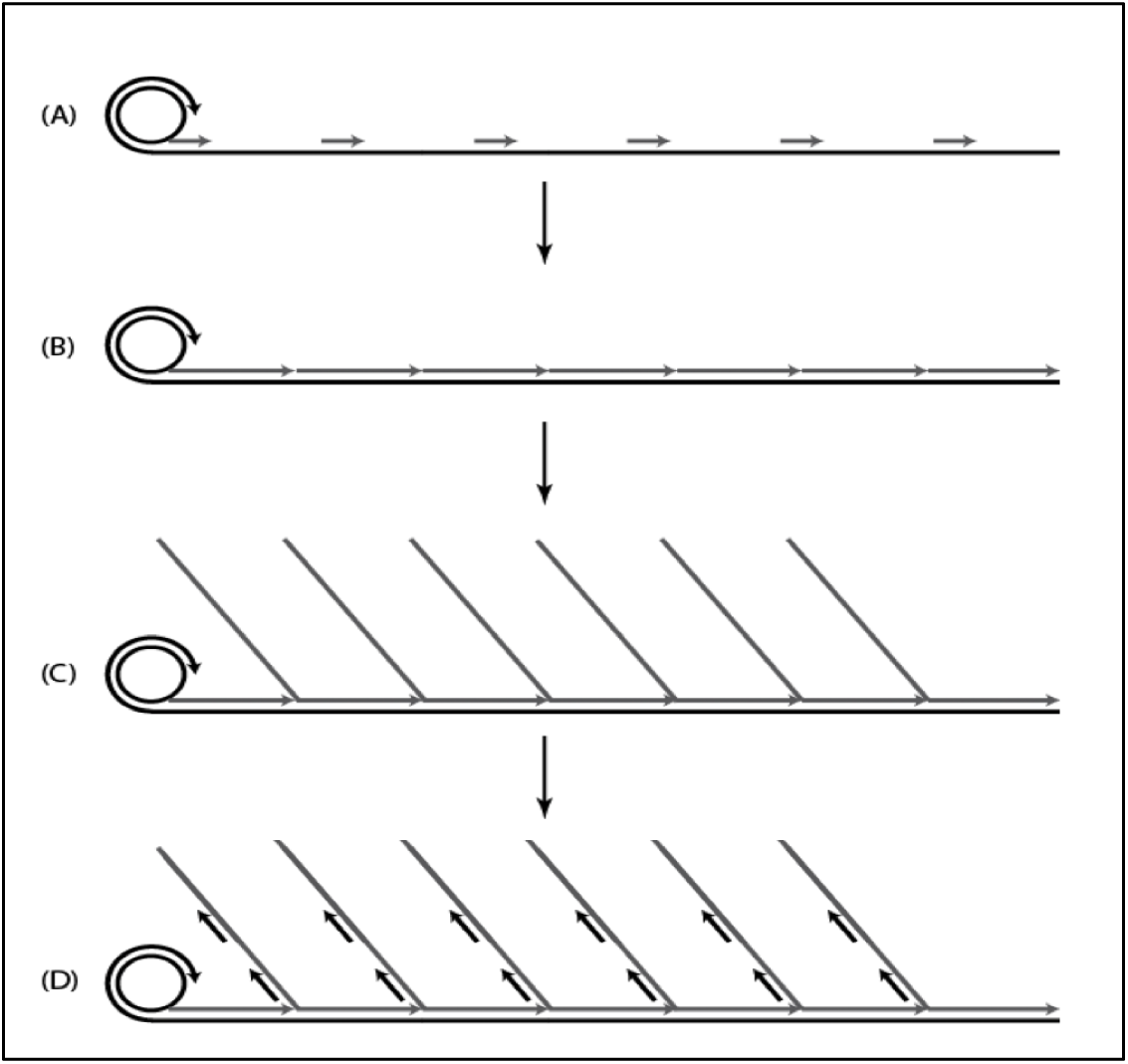
Illustration of complementary strand making and pair-end sequencing on the DNBSEQ MPS platform. (A) DNA nanoball (DNB), as a concatemer, containing copies of adaptor sequence and inserted genomic DNA, is hybridized with a primer for the first-end sequencing. (B) After generating the first-end read, controlled, continued extension is performed by a strand displacing DNA polymerase to generate a plurality of complementary strands. (C) When the 3’ ends of the newly synthesized strands reach the 5’ ends of the downstream strands, the 5’ ends are displaced by the DNA polymerase generating ssDNA overhangs creating a “branched DNB”. (D) A second-end sequencing primer is hybridized to the adaptor copies in the newly created branches to generate a second-end read. (C) and (D) are idealized drawings; not all branches are generated, and they are of different size.

MGI (a BGI subsidiary, Shenzhen, China) standard MPS kits were modified to implement the CoolMPS™ process. Labeled RTs were replaced by unlabeled RTs with a natural nucleobase (also obtained from MGI) and a cocktail of the four labeled antibodies (specific for each natural nucleobase) in binding buffer was added to the cartridge. The antibodies were labeled with fluorescent dyes of similar excitation and emission spectra as labeled RTs to enable imaging on the current sequencers. Antibody generation, selection, manufacturing, purification and labeling is described above in the antibody preparation section.

Each cycle of sequencing included reversible terminator incorporation with a modified polymerase, followed by binding of antibody. After washing excess, un-bound antibodies, standard imaging was performed, followed by bound antibody removal and standard 3’ de-blocking as either one combined, or two separate steps.

#### Antibody evaluation in sequencing assays on the DNBSEQ platform

In the initial DNBSEQ screening of several ELISA positive antibodies for each of the four nucleotides, we found that up to 50% had relatively weak positive signals. A possible explanation was unsuccessful clonal expansion or false positive ELISA. After several rounds of screening we finally selected a set of four antibodies with good signal and low background. We then evaluated properties of these antibodies required for sequencing. Primary splenocyte supernatant from promising clones was also screened by functional assay on the DNBSEQ platform using fluorescently labeled secondary antibody for specific signal and low residue levels.

##### Specificity

Accurate sequence determination requires that the antibodies are specific for the base associated with the 3’ reversible terminated ribose. To demonstrate that each antibody species is specific for each individual base, arrays of DNA nanoballs were created and hybridized with primers that were then extended one nucleotide with a reversible terminator. Fig. 4a shows the fluorescent intensity for populations of DNBs within a single imaging field after binding four labeled antibodies. Pairs of channels that do not have spectral dye crosstalk such as A-G, A-C, T-G, T-C do not show any antibody cross binding. DNBs are either negative in both channels or positive in one but not in the other channel (DNB clusters on the x and y axis). We attribute high specificity to our positive and negative antibody selection.

**Figure 4a:**
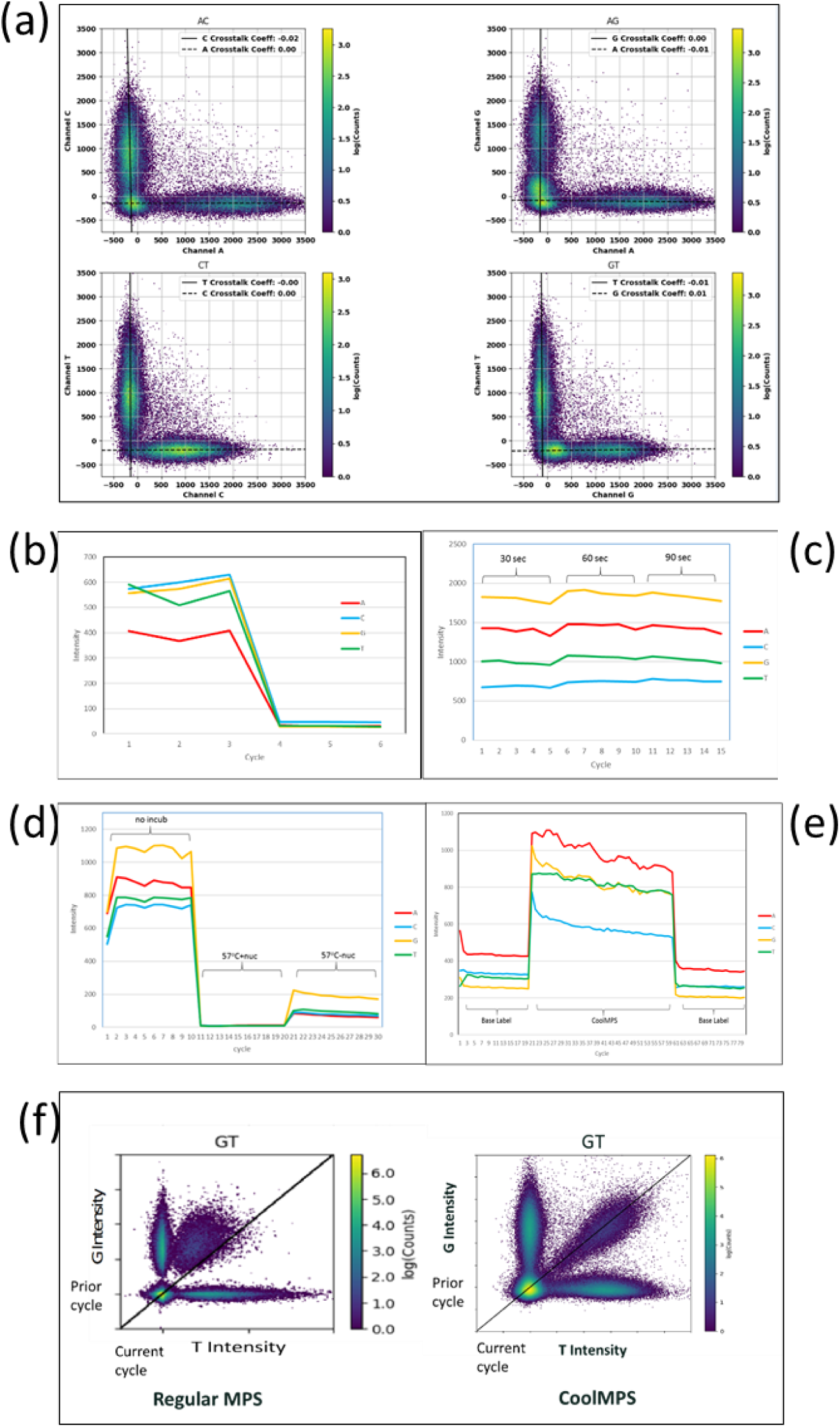
High base specificity of antibody binding. CoolMPS™ generated DNB intensities from one cycle are plotted in pairs of imaging channels. A random selection of 100,000 DNBs in an FOV are represented. Background subtracted intensities without dye crosstalk correction are presented. Only pairs of channels without dye cross talk are shown. For each pair, three clusters of DNBs are expected if there is no antibody cross binding on an X-Y co-ordinate representation: -/-; low X and Y intensities (clustered at zero co-ordinates), +/-; high X and low Y intensities (aligned with x axis), -/+; low X and high Y intensities (aligned with y axis). If there is cross-binding, +/- or -/+ clusters would shift from X or Y at an angle. In all four pairs, strong binding (relative signal in the range of 1000 counts) of only one antibody is observed without detectable cross-binding. **Figure 4b**: Antibody binding is dependent on both the base and the sugar with a 3’ –O-azidomethyl block. Three regular sequencing cycles in which the 3’ blocking group is removed after antibody binding and imaging, were followed by three cycles in which the 3’ blocking group was cleaved before antibody binding and imaging. Background subtracted, phase corrected and spectral cross-talk corrected intensities are shown and for each imaging channel (corresponding to each base) an average intensity of DNBs with highest intensities in that channel are depicted. **Figure 4c:** Antibody binding time. The same concentration of antibodies (∼4ug/ml, providing excess of antibodies) were allowed to bind to DNBs for either 30 sec, 60 sec or 90 sec at 35°C. 30 sec already generates >90% of maximal signal demonstrating fast binding kinetics of all 4 selected antibodies. Background subtracted, phase corrected and spectral cross-talk corrected intensities are shown and for each imaging channel (corresponding to each base) an average intensity of DNBs with highest intensities in that channel are depicted. **Figure 4d**: Removal of fluorescent antibodies after binding to RTs. In cycles 1-10 flow cells were washed briefly with pH7 SSC buffer at 40°C before imaging at 20°C. In cycles 11-20 flow cells were incubated at 57°C for 1 minute in 50mM Tris pH 9 buffer including RTs, for 60 sec before imaging. Cycles 21-30 show intensities after incubation for 60 seconds in the same buffer without nucleotides before imaging. Background subtracted and spectral cross-talk corrected intensities are used and for each imaging channel (corresponding to each base) an average intensity of DNBs with highest intensities in that channel are depicted. **Figure 4e:** Twenty cycles of base-labeled sequencing were performed before switching to CoolMPS™ sequencing (cycles 21-60), and then back to standard direct base labeled sequencing. Background subtracted, phase corrected and spectral cross-talk corrected intensities are shown for each imaging channel (corresponding to each base). An average intensity of DNBs with highest intensities in that channel are depicted for each cycle. **Figure 4f:** CoolMPS™ eliminates signal suppression. DNB signals in a set of DNBs are compared in channel G for the prior cycle (Y axes) and channel T for the current cycle (X axes). Labeled RTs chemistry and CoolMPS™ chemistry (natural unlabeled bases, labeled base-specific antibodies) are shown. Each point on the plot is a DNB forming 4 clusters: nonG/nonT, G/nonT, T/nonG and G/T. Lower than expected T signal is observed in the case of labeled RTs (the cluster of GT DNBs is shifted toward Y axes). No suppression was observed in CoolMPS™.

#### Binding requires the 3’ blocking group

Our immunogens use nucleotides with a 3’ -O-azidomethyl blocking group. After confirming base specificity (Fig. 4a) we next determined if the blocking group is required for strong binding. Fig. 4b shows in the first 3 cycles the intensity of fluorescence achieved when individual antibodies were incubated with the surface associated DNBs. Here, we report intensity as a background subtracted and spectral crosstalk corrected measure of the average population intensity for DNBs assigned to a fluorophore channel (having the strongest intensities in that channel) within an imaging field. All four antibodies produce strong signal (400-600 counts) when the blocking group was present during antibody binding. In cycle 4 onward, each cycle had a cleavage step before antibody binding. No signal detection was evident after removal of the 3’ blocking group suggesting that in addition to the base this chemical moiety is important for strong antibody binding potentially preventing antibody to bind to other target bases in DNA. Bases on non-terminal nucleotides can also be discriminate by other spatial or chemical features because they have a stacking base and phosphate on the 5’ and 3’ side.

##### Fast binding kinetics

In optimizing antibody-binding conditions we found that low salt (50mM) Tris buffer (pH7.6) provided efficient binding at 35-40°C. Fig. 4c shows the effect of 30, 60 or 90 seconds of labeled antibody binding to unlabeled RT nucleotides incorporated by DNBSEQ sequencing. Minimal increase in fluorescent intensity was observed with increasing times of incubation. Although this suggests shorter incubation time than 30 seconds is possible, it must be remembered that this represents the behavior of the population average and specific sequence contexts could behave differently. We attribute this fast binding to our double selection of strong binders, first by ELISA test and then in the initial screening using RTs incorporated by primer extension on DNB arrays. Furthermore, our optimized antibody-binding buffer is expected to enhance DNA-end breathing making the incorporated unlabeled RTs accessible to antibodies.

##### Efficient removal of bound antibodies

Complete removal of the bound antibodies after imaging and before the next cycle of nucleotide incorporation is important for high quality sequencing. It would be time beneficial if antibody removal and 3’ block cleavage could be done at the same time. We found that high pH (>pH8) and temperatures over 55 °C were efficient in quantitative antibody removal (Fig. 4d). We also found that including unlabeled RTs in the removal buffer speeds up the dissociation. Buffer conditions without including RTs are compatible with the cleavage reaction.

#### Labeled antibodies generate stronger signal than labeled RTs

Labeled RTs can have only one dye attached to a base due to proximity quenching. To minimize negative impact of base scar, usually only 60-70% are labeled. CoolMPS™ antibodies can be labeled with multiple dye molecules per antibody molecule potentially providing stronger sequencing signal. We tested the signal strength provided by the current random labeling process that balances the number of fluorophores per antibody and antibody inactivation.

Fig. 4e shows the relative intensities of base-labeled nucleotides over the first 20 cycle positions followed by an additional 40 cycle positions with antibody labeled detection, before returning to base-labeled RTs. Relative to base-labeling of nucleotides, antibody detection generated much stronger signal with some fluorophores producing an over 300% increase in intensity relative to its base-labeled counterpart. The range of responses by different fluorophores may reflect labeling efficiency of the dyes to the specific antibodies, antibody binding affinities, or fluorophore quenching. The benefits of increased intensity include preservation of sufficient signal in low template copy DNBs in high density nanoarrays throughout long sequencing runs, less intensive exposure or more rapid imaging.

#### No signal suppression

We observed that labeled RTs generate some signal suppression (e.g. quenching) in the following cycle which most likely was due to modified (“scarred”) bases. Because CoolMPS™ uses unlabeled RTs we expected no such effect. Fig. 4f compares signals in a set of DNBs (from one field-of-view) in two consecutive cycles and demonstrates that DNBs that have G at the prior cycle and T in the current cycle have a suppressed T signal when labeled RTs were used. Lower than expected T signal causes the GT cluster to move from the diagonal toward the Y axis, representing G signals. No suppression was observed in CoolMPS™ using unlabeled RTs with a natural base without any scar. Furthermore, dyes on the T antibody are further from the G base avoiding quenching. This is one of many advantages of CoolMPS™ (Table 1, discussion).

**Table 1:** Estimated sequencing error rates in high quality DNBs (a single DNB with sufficient template copies per site) for 4-color sequencing on 2-color imager (4cs2ci) and standard 4-color sequencing on 4-color imager (4cs4ci). Discordances in base calls with quality scores less than Q20 are treated as sequencing errors and others as true sample polymorphisms or library (e.g. PCR) errors. Four specific error types (A/T, C/G, G/C, T/A) affected by dye-crosstalk in 4cs4ci are also shown.

|  |  | 4cs4ci |  | 4cs2ci |  | Error Reduction (fold) |
| --- | --- | --- | --- | --- | --- | --- |
| Read Yield after filtering (%) |  | 74.2 |  | 79.8 |  |  |
| Reference base | Called base | Color | Error rate (%) | Color | Error rate (%) |  |
| A | non A | 1 | 0.00533 | 1 | 0.00150 | 3.6 |
| C | non C | 4 | 0.00257 | 2 | 0.00045 | 5.7 |
| G | non G | 2 | 0.01313 | 2 | 0.00070 | 18.8 |
| T | non T | 3 | 0.00373 | 1 | 0.00225 | 1.7 |
| All bases | Non-reference |  | 0.00619 |  | 0.00123 | 5.1 |
| A | T |  | 0.00167 |  | 0.00050 | 3.3 |
| C | G |  | 0.00093 |  | 0.00010 | 9.3 |
| G | C |  | 0.01043 |  | 0.00020 | 52.2 |
| T | A |  | 0.00130 |  | 0.00110 | 1.2 |

### Full sequencing tests of CoolMPS™ chemistry

#### Generating 200 base reads: SE200 sequencing

MPS reads longer that 100 bases are very useful. As an initial demonstration test of CoolMPS™ potential we obtained 200-base reads. Two hundred cycles of sequencing were performed on DNBs loaded into the lanes of a flow cell of a DNBSEQ-G400 sequencer. DNBs were prepared from standard 300-base libraries of E. coli DNA using MGI’s protocols. Fig. 5a shows the average called-base intensity of DNBs in a selected region of the array with optimal fluidics and optics to highlight potential of this new chemistry.

**Figure 5a:**
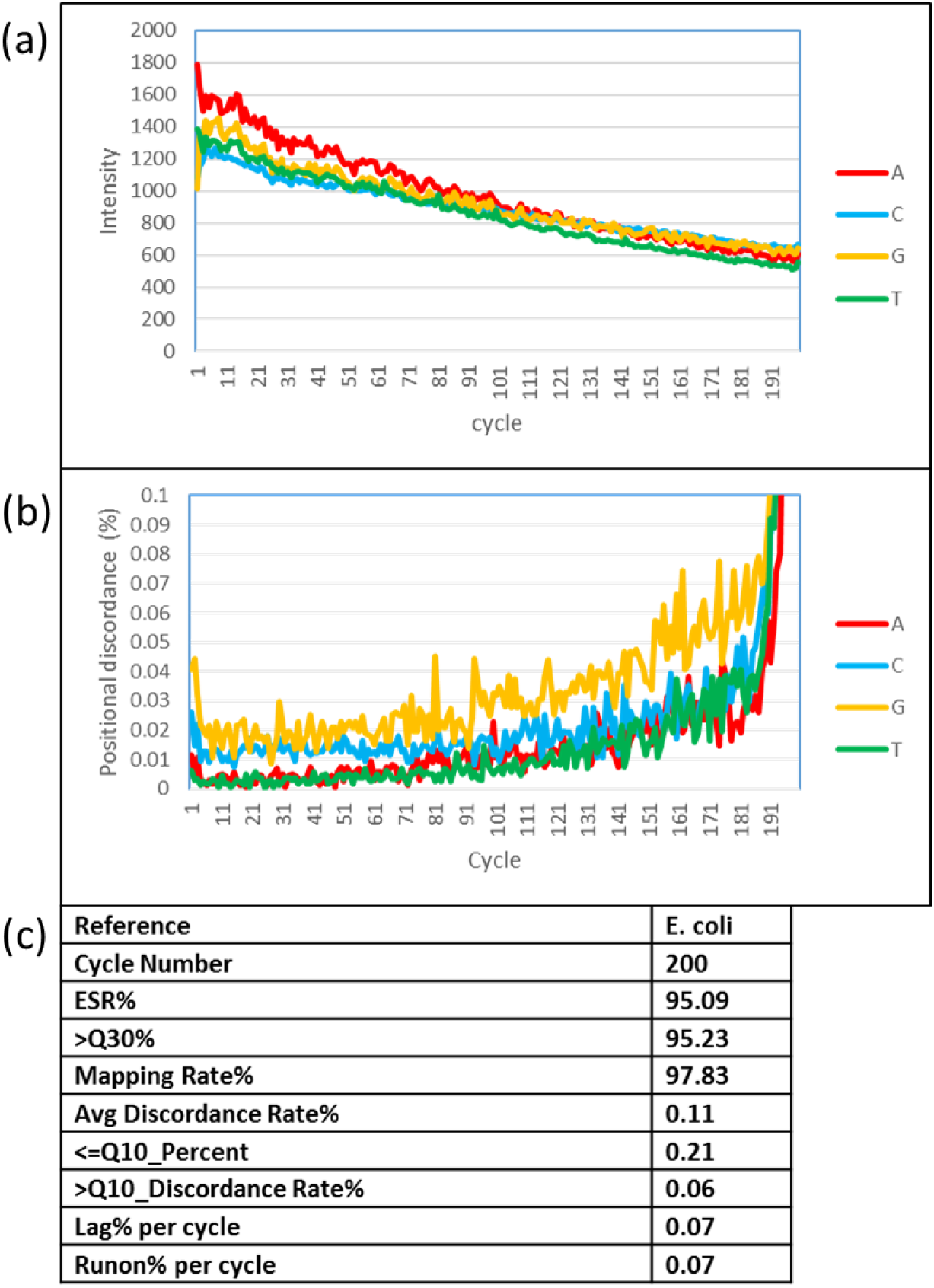
Intensity change over 200 cycles of single-end read. Background subtracted, phase corrected and spectral cross-talk corrected intensities are shown and for each imaging channel (corresponding to each base) an average intensity of DNBs with highest intensities in that channel are depicted. **Figure 5b:** Positional discordance for 200 cycles of SE sequencing. Note; the high rate of discordance increase after cycle 185 is due to short inserts and reading into the adapter region not matching the human reference. **Figure 5c:** Key performance metrics for SE200.

As previously observed with directly labeled nucleotides, a decline of intensity was observed as cycles progressed. Several factors contribute to this including i) out-of-phase signal, ii) irreversible termination, in part due to impure RTs and DNA damage in imaging, or ii) DNA loss. We found similar signal loss even without antibody binding and imaging, just cycles of incorporation of unlabeled RTs (data not shown). This excludes the impact of antibody binding, imaging or removal. Differences in decline rates between the bases are presumed to be due to the influence of changing background or efficiency of illumination of light collection during cycles. Although declines in dye intensity could be occurring in the cartridge during the run, this is a minor contributor since minimal increase in intensity was observed when fresh reagents were added through the course of a run (data not shown). Nevertheless, the remaining signal after 200 cycles is still high, supporting the possibility of much longer CoolMPS™ reads.

Positional discordance is increasing over cycles as in the standard MPS with reversible terminators. This is due to i) accumulation of out-phase signal that becomes confused with dye-cross talk and ii) signal loss relative to background, especially affecting DNBs with low template copy number. Lag (−1 signal) and runon (+1 signal) were relatively low per cycle (<0.1%) but still accumulated to ∼30% combined out-of-phase in 200 cycles. As shown in Fig. 5c after filtering out 5% of empty spots and spots with two or more DNBs from all binding spots in the array, the mapping rate of the remaining 95% of DNBs was 97.83% with an overall discordance of 0.11% which was further reduced to 0.06% in 99.79% of base calls with a quality score >Q10. This is a very promising result for 200 base reads showing high accuracy and 93% sequencing yield (0.95 filtered reads x 97.83% mapping rate).

We further evaluated sequence discordance in 100-base reads in a PCR free E. coli library, avoiding all PCR errors because DNBSEQ also does not use PCR to clonally amplify DNA for sequencing. We used DNBs in a selected region of the array with optimal fluidics and optics and filtered out mixed (two DNBs per spot) and small DNBs. We obtained overall discordance of 0.029% (1 difference from the reference in 3,500 called bases). We then calculated discordance at different base-call quality filters. Base calls with quality score >20 (close to 99.8% of all base calls) have five to six-fold less errors (discordance close to 0.005% or one mismatch in 20,000 bases). The remaining high-quality discordances can be caused by replication errors in DNA, DNA damage or real sequencing errors. This indicates great potential of CoolMPS™ for high-quality sequencing with very low overall error rate and extremely rare sequencing errors with high-quality base-calls.

#### High CoolMPS™ signal improves sequence quality in DNBs with a low template copy number

As shown in Fig. 4e, CoolMPS™ chemistry provides an approximately three-fold higher signal than standard MPS on the same DNB array. We evaluated the benefit of such high signal for sequencing DNBs with a small number of template copies (Fig. 6a). This is important for sequencing long templates (e.g. ∼1000b) using small DNBs in high density nanoarrays that are expected to have less than 100 template copies. Fig. 6b shows that DNBs from a human genomic library with less than 50 copies of 400 base average template size (prepared by a 10-min RCR reaction) can be sequenced with CoolMPS™ with a low estimated error rate of 0.055%, 3-fold lower than that obtained in standard MPS with dye-labeled RTs. This clearly demonstrates the benefits of 3-fold stronger signal obtained by antibodies labeled with multiple dye molecules. The benefit of CoolMPS™ is still significant, but lower, for DNBs with more than 100 copies (e.g. 25 min DNBs).

**Figure 6(a):**
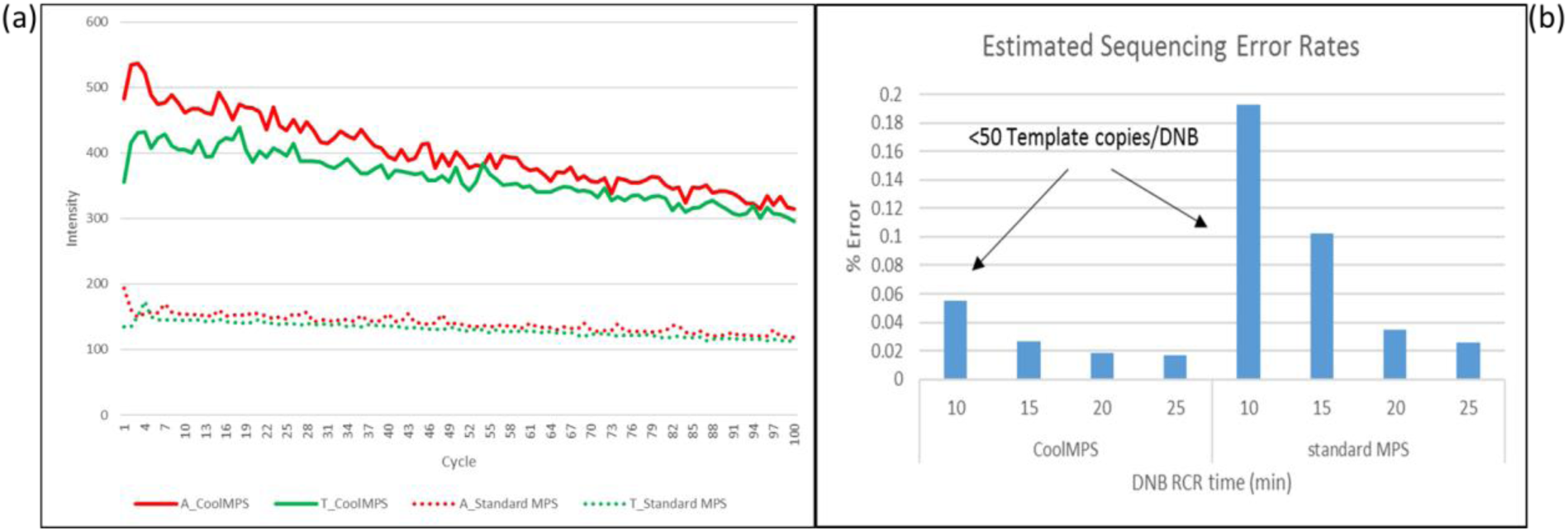
A and T channel intensities over 100 cycles for DNBs generated by 10 min RCR time and sequenced by standard MPS or CoolMPS™. A 3-fold higher signal is observed with CoolMPS™. **Figure 6(b)**: Error rates for DNBs of q-scores less than 20 (q<20) showing that standard chemistry is less accurate than CoolMPS, especially for smaller DNBs.

#### High quality 150-base pair-end reads: PE150 sequencing

Pair-end (PE) sequencing provides very useful MPS reads that bridge repeats longer than reads and minimizes the need for long continuous reads. PE150 (150 bases from both ends of 300-600b inserts) is most frequently used.

We tested CoolMPS™ PE150 to demonstrate that using antibodies does not interfere with the DNBSEQ PE process of controlled MDA depicted in Fig. 3. Figure 7(a) shows the change in intensity over the 150 cycles of the first strand, then good recovery of intensity on the second strand as the complementary template and corresponding sequencing primer was used for extension. In this test, the concentration of antibodies used for the second strand was twice that of the first strand. Overall there was about a 30-50 % decline in intensity values over the 150 positions of the first strand and a 40-50% decline on the second strand, in part due to higher incorporation incompletion (lag) in the second strand. After filtering about 11-13% of empty and low-quality array spots, mapping rates were >99% with a discordance rate of 0.08% and 0.26% on the first strand of E. coli (300b inserts) and Human (400b inserts) DNA libraries, respectively (Fig. 7(b)). For the second strand, the mapping rate was about 99% with a discordance rate of 0.22% and 0.62%, respectively. After filtering 0.4% and 0.8% of base calls with quality score <10, the combined discordance was reduced to 0.06% and 0.24% respectively in E.coli and Human DNA libraries. Part of discordance rates were due to PCR errors introduced in library preparation and human libraries were expected to have higher discordance compared with E.coli libraries. This is due to more polymorphisms in the human sample relative to the human reference compared with the E.coli sample relative to the E.coli reference.

**Figure 7a:**
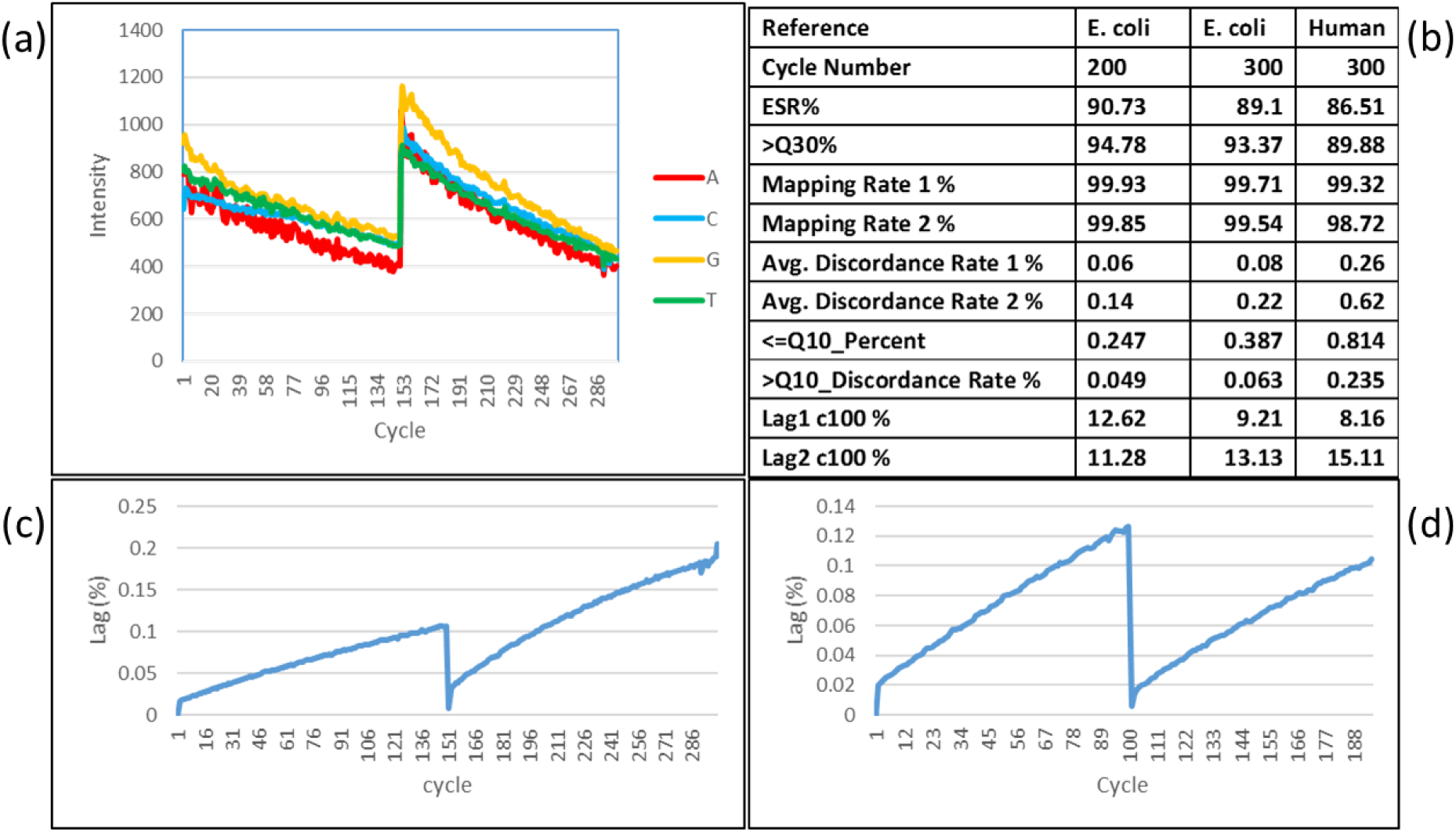
PE150 intensity for a Human DNA library. Background subtracted and spectral cross-talk corrected intensities are shown, and for each imaging channel (corresponding to each base) an average intensity of DNBs with highest intensities in that channel are depicted. **Figure 7b:** Key performance metrics for PE150. **Figure 7c:** PE150 Lag (−1 out-of-phase incorporation) in the run shown on Fig. 6a. Lag represents intensity contributions of the prior (−1) base to the current cycle. **Figure 7d:** PE100 Lag in a PE100 run (E. coli library) with optimized Phi29 removal.

In spite of a higher signal, the higher discordance rate in the second read was due to higher lag and lower quality thresholds used for DNB filtering. Higher lag (−1 out-of-phase) in the second read is probably due to incomplete removal of Phi 29 polymerase used for the complementary strand making. This was confirmed in a PE100 run using optimized Phi29 removal, reducing accumulated lag from about 15% to about 11% (Figures 7(b), 7(c), 7(d)). Furthermore, the lag accumulation is more linear indicating less of -2 phase. This result illustrates the complexities (many biochemical steps with multiple interdependences) of MPS process that require carefully balanced optimizations.

Overall these results demonstrate that CoolMPS™ chemistry is ready for popular PE150 sequencing, although still at the beginning of the development cycle.

##### Demonstrating potential of CoolMPS™ for longer MPS reads

As a demonstration of longer MPS read potential of CoolMPS™ we obtained single-end 400 base reads on a human library and standard DNB-nanoarrays (less than 100 copies per DNB of a 400- to 600-base genomic template). The distribution of q-scores across quality bins over 400 bases in the array regions with minimal optical and fluidic aberrations (to focus on CoolMPS chemistry performance) are shown in Figure 8. The average Q30 fraction for 400 bases was 91%. At a 75% filtered read yield, an acceptable sequencing error rate of 0.42% was estimated in the first 390 bases (avoiding the last 10 bases that may contain adapter sequence). Two-thirds of these errors (0.27/0.42=64%) were found in a small fraction (0.84%) of very low quality (<Q10) scores and could be filtered out by conversion to no-calls (3-4 no-calls per 400 base read, on average), reducing the error-rate to 0.15%. The error rate in the first 300 bases was 0.14% indicating that 67% (0.42-0.14/0.42) of the errors in the 390 base-reads were in the last 90 bases. This is expected due to accumulation of out-of-phase signal, combined with signal stochastic on <30 template copies usable after 300b in this proof-of-concept demonstration of long CoolMPS™ reads. A further increase of template copies per DNB and dyes per antibody, combined with a further reduction of out-of-phase incorporation and signal loss, is expected to produce higher quality of even longer CoolMPS™ reads matching the read length of Sanger sequencing (e.g. 500-700 bases).

**Figure 8:**
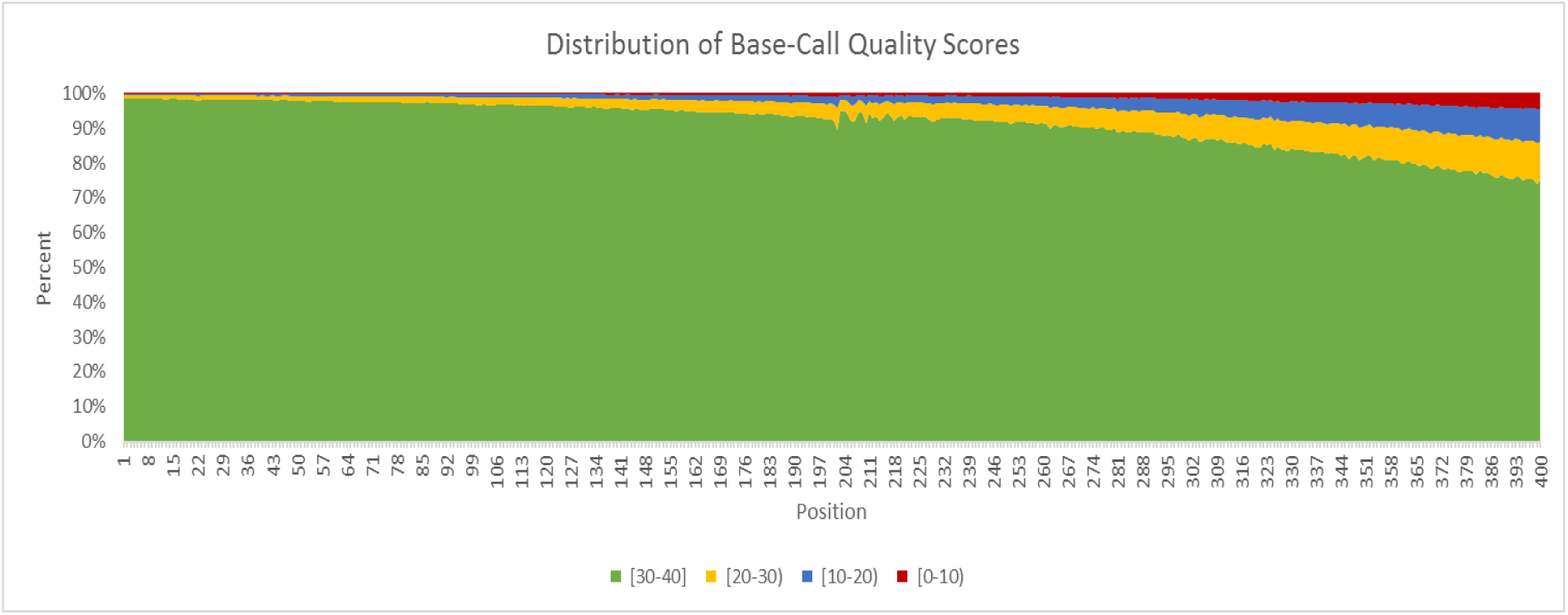
Distribution of base-call Q-scores by position over 400 sequencing cycles. For the last 200 bases, two-times higher antibody concentration was used.

#### “Four-color” CoolMPS™ sequencing on two-color imagers: The most accurate MPS reads

In standard 4-color sequencing, 4 fluorophores with distinct colors are typically assigned to the 4 different bases. One limitation of this approach is the spectral overlap of many commonly used dyes and the requirement for narrow band-pass filters to minimize the level of such dye cross-talk. These band pass filters, in addition to adding complexity and cost, also restrict the amount of light that can be collected from each fluorophore. To overcome some of these challenges 2-color sequencing has been developed (*16*) which utilizes just two fluorophores that are spectrally more distantly separated to identify the four bases from two images. One drawback of this technology however is that one of the bases is typically assigned a null intensity value and one of the bases is assigned a mixture of the two colors. This can still result in overlap of base-call clusters, particularly for weaker intensities that can make base-calling challenging.

CoolMPS™ can solve many of these issues by utilizing two-color imaging systems with their inherently clean and strong fluorescent signals as well as lower costs to obtain four distinct images (one for each base) in two consecutive binding/reading/removal steps of two fluorescently labeled antibodies at a time during each sequencing cycle. We term this method “4-color sequencing” on 2-color imagers (4CS2CI)

To assess the sequencing quality using such an approach we performed 100 cycles of single-end sequencing using a DNBSEQ-G50 instrument which is based on two color excitation and emission. After incorporating all 4 unlabeled RTs, In the first stage of detection, only antibodies for two bases, A and G, labeled with dye-1 and dye-2 respectively were allowed to bind. After imaging, the antibodies were quickly removed in an optimized displacement reaction. Antibodies for T and C were then allowed to bind, again each labeled with dye-1 and dye-2 respectively, before imaging once more to obtain a total of four distinct images without dye-crosstalk, one for each base. Differential detection in this way effectively eliminated spectral crosstalk and allowed each called base to be represented with a single, positive, intensity value. Furthermore, the dye with longest wavelength in regular 4-color sequencing is not used, reducing light bleed between neighboring DNBs.

Table 1. shows the discordance for DNBs at 80% of DNB binding sites (80% read yield) with high quality DNBs (a single DNB with sufficient copies per site) and base calls with quality scores less than Q20. This sub-population of discordances is more representative of sequencing errors by minimizing those discordances introduced from library preparation (e.g. by PCR) or true sample polymorphisms that are expected to have high quality scores, over Q30, on average.

As expected, the results show near elimination of dye cross-talk and long-wave length light bleed related G to C errors (52.2 fold G to C error reduction), leading to an exceptionally low average error rate for C (0.00045%) and G (0.00070%) bases in raw reads. This is 1 error in 170kb (0.00058%), on average, an order of magnitude lower than in standard 4CS4CI CoolMPS™. A and T bases labeled with dye emitting color-1 show less error reduction due to non-uniform illumination of this dye on the 2-color imager used for these experiments. If 0.0064% of bases called with Q<10 are converted to no-calls (1 no-call in 15kb) the remaining errors in these 100-base raw reads would be close to 1 in 1Mb. Thus, the CoolMPS™ enabled 4CS2CI method is promising to provide the most accurate and efficient MPS sequencing.

## Discussion

We have demonstrated, for the first time, sequencing of DNA utilizing the specificity of natural nucleobase recognition by fluorescently labeled antibodies including accurate SE400 and PE150. This novel methodology, is not only rapid enough to compete with existing commercially available MPS methods based on fluorescent dyes covalently linked to bases, but it also offers high accuracy (e.g. 1 error in 20,000 raw base calls with quality score >20 in good size DNBs) and potentially longer reads and lower cost at scale. There are other methods that can detect unlabeled RTs (*17*) but they are limited to incorporation of one nucleotide at a time. Having four base-specific antibodies allows the incorporation of all four unlabeled RTs to arrayed DNA in one reaction providing speed and minimizing false nucleotide incorporation. Furthermore, antibodies are broadly used in diagnostic tests and as therapeutics. There are many developed tools to further optimize the CoolMPS™ process, including replacing full antibodies with smaller versions such as ScFv or nanobodies expressed in bacterial host and efficiently labeled at targeted sites.

DNA base recognition by antibodies has been described in the past, usually for the detection of chemically modified nucleotides (*18*–*20*). The monoclonal antibodies described in this report not only recognize the natural base type (whether it be A, C, G or T) but also bind to a small reversible blocking group at the 3’ end of the nucleotide. Others reported discriminative binding of an antibody to a modified G with 2’ methyl group versus a similarly modified A, with a preferential binding to the base and additional binding to the modified sugar (*21*). The binding was tested using free nucleotides in a competitive binding assay. Our results demonstrate that all four 3’ sugar-modified nucleotides (four 3’ –O-azidomethyl RTs with natural base) can be efficiently discriminated in the context of dsDNA after incorporation of RTs by polymerase at the end of an extending strand hybridized to a longer overhanging template (Fig. 4a). The amazing specificity of these antibodies discriminates C RT from T RT in spite of sharing the 3’ blocking group required for antibody binding (Fig. 4b) and having just a small chemical change on the single ring nucleobase. Similarly, full discrimination is achieved for A and G RTs, both with two ring nucleobases. Over 80% of monoclonal antibodies entering negative selection have good base specificity. Furthermore, selected antibodies do not bind 2’ -O-azidomethyl nucleotides (obtained from Jena Biosciences, Germany) even though the same polymerase incorporates 3’ and 2’ -O-azidomethyl RTs. This strong discriminative binding to the terminal RT incorporated in one strand of dsDNA enables the use of these antibodies for accurate base reading in a MPS process.

Binding times of antibodies were demonstrated to be relatively quick compared to many common procedures utilizing antibodies for detection (eg western blot, ELISA) with just 30 seconds proving effective for generating enough intensity to provide low-error base-calling. Increased antibody binding time had minimal effect on increasing intensity, suggesting most available target sites were occupied within 30 seconds. Furthermore, about 4ug/ml of antibodies is enough to bind most of incorporated RTs. This is surprising because the target nucleotide is present in dsDNA and the immunogen used was single monophosphate reversibly terminated nucleotide. Most likely there is some temporary dsDNA end-melting allowing antibody to bind. Preferred binding buffer with low salt and no Mg++ (that helps breathing of DNA ends) supports this explanation.

Using unlabeled RTs and labeled antibodies specific for each natural nucleobase for massively parallel sequencing provides multiple process advantages leading to higher accuracy, longer reads, higher throughput, lower cost and other performance benefits (Table 2).

**Table 2:** *CoolMPS™ advantages*

| Cool MPS Features | Process Benefits | Performance Benefits |
| --- | --- | --- |
| Unlabeled RTs used for sequencing cycles | Less expensive to make | Lower cost |
|  | Easier to incorporate to completion (less out of phase) | Higher Quality<br>Longer Read |
|  | Provide natural bases after deblocking RT without interference with the next sequencing cycle and a more stable dsDNA helix | High Quality<br>Longer Read |
| Base specific labeled antibodies used for base reading | Multiple dyes per one Ab provides a) accurate sequencing of DNBs with a small number of template copies in dense nanoarrays or long templates, and b) brighter signals for more efficient imaging with simpler imagers | High Quality<br>Longer Read<br>Lower Cost |
|  | Less DNA damage (dyes are further from DNA) | High Quality<br>Longer Read |
|  | Flexibility in balancing signals, e.g. in 2-color MPS using only two dyes | High Quality<br>Low cost |
|  | Stepwise base detection of 4 bases in 2- or 1-color simpler and more efficient imagers (e.g. "4-color" sequencing on 2-color imager) | Higher Quality<br>Lower cost |
| Implemented on DNBSEQ MPS platform | PCR-free, error-free DNA nanoarrays without barcode swapping | Higher Quality Higher Sensitivity |
|  | High Array Density enabled by 200nm DNB size | Lower Cost<br>Higher Throughput |
|  | High DNA density per spot, easier to image, higher SNR. | Higher Quality<br>Lower Cost |
|  | Over 90% of DNBs made are loaded without overloading | Lower DNA input |

First, unlabeled RTs are more efficient to incorporate. For example, using a modified polymerase (BGI Research, Shenzhen, China) on the DNBSEQ platform we measured almost 3 times more efficient incorporation of unlabeled RTs resulting in three to four times lower out-of-phase due to incompletion of incorporation (0.26% unlabeled vs. 0.87% labeled per cycle for 1min incorporation at 1μM nucleotide concentration).

Because no dye is linked to the nucleotide, there is no residual scar after cleavage of the dye. Since the 3’ block is also converted back to a free hydroxyl group the resultant nucleotide structure is identical to a natural nucleotide at the end of each sequencing cycle. Residual cleavage scars could affect the flexibility of the newly synthesized strand and the binding efficiency of polymerase for continued extension of the strand or suppress the signal in the following cycle as shown in Fig. 4f. Nucleotide synthesis costs are also dramatically increased when cleavable dyes are added to the nucleotide and high purity is usually required because of possible termination by-product reactions that can occur during the nucleotide labeling procedure. Termination by-products have a detrimental effect on longer read sequencing due to the accumulated loss of intensity at each target site. Use of labeled antibodies that recognize a non-dye label (e.g. biotin attached to the base) has been considered for sequencing for a long time but issues such as base-scar and cost of synthesis of these labeled RTs remain. Furthermore, attaching labels to the 3’ block impacts incorporation due to total size of these 3’ modifications.

Our new method utilizes a standard labeling methodology in which 10-100 micromolar of NHS ester (succinimidyl ester) modified dye is reacted with 1 mg/ml protein. Even without further purification of free unreacted dye from antibody, this results in final dye concentration in the antibody binding reaction of 0.2 μM compared with, in excess of 1 μM typically used of highly purified base-labeled labeled nucleotides. In addition to very standardized and developed labeling methodologies for antibodies, the availability of in-vitro produced, and high-scale quantities of, monoclonal antibodies is now routine and at low cost (*22, 23*).

Attaching multiple dye molecules per antibody provides stronger signal than one dye molecule attached to the base for more efficient high-quality imaging with less illumination light. Due to much stronger signals we show high quality CoolMPS™ sequencing of DNBs with less than 50 template copies (Fig. 6). This opens the possibility of MPS on ultra-high density DNB nanoarrays comprising small DNBs (e.g. <100nm). This also allows to balance signal intensities in 2-color MPS sequencing proposed in 2007 (*16*) where one nucleotide has to be detected at two distinct wavelength channels. More dye molecules can be attached to the antibody where 50% of antibody molecules have to be labeled with one dye and 50% with a different dye.

A special benefit of CoolMPS™ is the possibility of stepwise base detection after single incorporation reaction of all unlabeled RTs. This is enabled by fast binding and removal of labeled antibodies without removing the 3’ blocking group. Each base can be detected in a separate image using more efficient and cost-effective 2- or 1-color imagers without dye crosstalk present in 4-color imagers. For 2-color imagers, two antibodies labeled with different dyes would be bound first and two images generated. After a quick removal of bound antibodies (possibly in less than 10 sec at >60°C in optimized displacement buffer), two other antibodies labeled with the same pair of dyes would be bound to generate two more images, one for each base. For the fast imagers, the entire “4-color sequencing” on 2-color imagers (4CS2CI) process will take slightly longer, but the sequence quality is expected to be much higher because 2-color imagers collect 2-3 more light (wider filter band) without any dye cross talk. In a proof-of-concept 4CS2CI demonstration we show an order of magnitude lower sequencing error rate (1 error in 170kb) primarily due to elimination of dye-cross-talk errors. With expected improvements in speed of imaging and increased need for maximally accurate clinical sequencing 4CS2CI CoolMPS™ may become the MPS method of choice.

Limitations of using antibodies for sequencing include high cost of set up, but once antibody clones are produced ongoing generation costs can be minimal. Additional time is required for the binding of antibodies, but this has shown to be as little as 30 seconds of additional time per cycle. Further optimization of CoolMPS™ antibodies, composition of binding reaction and speed of fluidics potentially can cut the antibody binding time to negligible levels.

CoolMPS™ technology is at the beginning of its development cycle and many future improvements are expected such as reduction of signal loss and out-of-phase reading, even brighter labeling, and further reduction of remaining low-quality base calls caused in part by sporadic partial loss of true signal. One of the CoolMPS™ chemistry goals is to utilize better incorporation of unlabeled RTs and stronger signal of labeled antibodies to achieve high-quality longer reads (e.g. 500+ bases). Full implementation of highly accurate “4-color” sequencing on efficient 2-color or 1-color imagers is another exciting future development.

In addition to PCR-free DNBSEQ MPS platform, CoolMPS™ can be used on any MPS platform, including PCR-based clonal arrays (PCR clusters on beads or directly on slides) or single molecule array. The combination of higher quality and lower cost of CoolMPS™ chemistry and PCR-free cost-effective DNB nanoarrays creates a novel advanced MPS platform to drive implementation of genomics-based health monitoring that requires comprehensive, accurate and affordable sequencing-based pre-symptomatic screening tests.

## Acknowledgments

We would like to thank Lingling Peng, Ann Tran, Kate Nguyen, Parul Agarwal, Hannah Debaets, Hongyu Chen and Puzhou Wang for technical support, Lin Wang and Tam Berntsen for libraries preparation, Sophie Liu for helping with drawing Fig. 1, and Naibo Yang for helpful discussions.

This work was supported by the Shenzhen Municipal Government of China Peacock Plan (No. KQTD2015033017150531).

